# Coevolution of structural variation and optimization in sound systems of human language

**DOI:** 10.1101/346965

**Authors:** Meng-Han Zhang, Tao Gong

## Abstract

Evolution of sound systems of human language is an optimization process to improve communicative effectiveness: The intrinsic structures of sound systems are constantly organized with respect to constraints in speech production and perception. However, there lack sufficient quantitative descriptions of this process, large-scale investigations on universal tendencies in sound systems, and explicit evidence on whether demographic and/or geographic factors can influence linguistic typology of sound systems. Here, we proposed two composite parameters, namely structural variation and optimization, to capture linguistic typology of sound systems, vowel systems in particular. Synchronic comparisons based on a large-scale vowel corpus of world languages revealed a universal negative correlation between the two parameters. Phylogenetic comparative analyses identified a correlated evolution of the two, but with distinct evolutionary modes: a gradual evolution of the structural variation and a punctuated equilibrium of the optimization. Mixed-effect models also reported significant effects of speaker population size and longitude on shaping vowel systems of world languages. All these findings elaborate the intrinsic evolutionary mechanism of sound systems, clarify the extrinsic, non-linguistic effects on shaping human sound systems, and quantitatively describe and interpret the evolution and typology of sound systems.

## Introduction

Research in linguistic typology attempts to not only specify language diversity but also generalize universals. Cross-linguistic surveys on grammatical (1, 2) and morphological systems (3, 4) of world languages have revealed many universal tendencies that are tied closely to human cognition and perception. However, there lack sufficient attentions to finding universal tendencies of sound systems. Among various components, sound systems are the most concrete and verifiable components of languages (5, 6). They exhibit great diversity in the numbers of consonants (ranging from 6, as in Rotokas, to 130, as in !XOO, with a mean of 22.8) and vowels (between 3, as in Aleut and 40, as in Dan, with a mean of 8.7) (7, 8), and undergo constant evolution driven by both intrinsic and extrinsic factors.

The functional goal of improving communicative effectiveness between speakers and listeners is one of the dominant intrinsic factors driving the adaptive self-organization of sound systems (9). Primary functional pressures for the evolution of sound systems include: perceptual dispersion (10), perceptual salience (11), and articulatory costs (12-14).

The dispersion theory states that the sound systems of human language are shaped predominantly by perceptual distinctiveness between sounds (10, 15). Taking the example of vowel systems, this theory defines *dispersion* as a systematic property of vowel systems measuring perceptual distinctions between vowels, and suggests that languages prefer systems in which vowels are distinct maximally from each other in a perceptual domain (7). The dispersion-focalization theory integrates *focalization* that reflects perceptual salience of vowels, and uses a weighted sum of dispersion and focalization to capture the global structural variation of vowel systems (11). Derived from the Quantal theory (13, 14), the concept of perceptual salience of vowels resembles that of focal colors (16, 17), and can be approximated by the first two formants of vowels (18, 19). Moreover, sound systems are also constrained by speech production. Articulatory costs such as *articulatory space area* appear evident in determining the structural plasticity of sound systems (20, 21). According to the articulatory complexity theory (21), the auditory pressures of maximizing perceptual distance and minimizing articulatory effort collectively shape the articulatory subspace of a sound system. From a phonological perspective, the Quantal theory (14) proposes that speech sounds easier or more reliable to produce appear more common in world languages, whereas those harder to produce are less so, which is supported by another theory of sound change (22). This offers an alternative way to characterize articulatory difficulty in speech production. In summary, dispersion, focalization and articulatory costs are three informative parameters capturing systematic properties of sound systems, vowel systems in particular. Due to functional pressures and self-organization, the sound systems of human language exhibit some universal characteristics.

In addition to intrinsic factors, whether and how extrinsic, non-linguistic factors affect the sound systems of human language has also been studied, but the conclusions remain controversial. For example, some studies discovered that the population size of a language casts a significant effect on the structure of its sound system (23-27), but other studies (28, 29) did not find explicit correlations between population size and sound system. Correlations between variations in sound systems and geographic distributions of languages seem evident in some study (23), but not in the other (30). Ongoing discussions on the ecological adaption of language are also subject to suspicion (31-33). All these indicate that whether non-linguistic factors contribute to language evolution still deserves a close inspection.

From an evolutionary perspective, the adaptive self-organization of sound systems, especially vowel systems, is practically viewed as an optimization process (9). However, there lack sufficient quantitative descriptions of this process, except for an unpublished preliminary model (34). As an important component of sound system, vowel system is a simple entry point into the study of language typology and speech evolution. In this paper, we thus examined the sound systems of human language based on Becker-Kristal’s vowel inventory corpus, which involves 532 language samples from 357 languages and dialects. We integrated the parameters of dispersion, focalization, and articulatory space to capture the structural variation of vowel systems, and designed a stochastic sampling approach with consideration of perceptual constraint to address the structural optimization of vowel systems. Synchronic comparisons of these parameters across languages revealed a universal correlation between the structural variation and optimization of vowel systems. Inter and intra-regional comparisons confirmed this universal tendency of vowel systems throughout different geographic regions of the world. From a diachronic perspective, we applied phylogenetic approaches on 33 contemporary Indo-European languages to identify whether the structural variation and optimization of vowel systems could coevolve and in what modes they evolve. Finally, we established mixed-effect models to clarify whether speaker population size and geographic distribution of languages are correlated with vowel systems. This study provides quantitative description and interpretation of the evolution of sound systems in human language.

## Results

### Principal component analysis (PCA) of the structural variation and optimization of vowel systems

To measure structural optimization of vowel systems, we applied the MCMC sampling procedure respectively on the three systematic properties, namely dispersion, focalization and articulatory space. The sampling procedure was modified from (34) with consideration of perceptual limitations (see Supplementary Information). After obtaining the optimization values of these systematic properties, unlike the summation function as in (11), we performed PCA separately to calculate the structural variation and optimization of vowel systems. As for variation, the first principal component of the three systematic properties could explain 98.48% of total variance. As for optimization, the first principal component of the three optimization values could explain 77.73% of total variance. There is a significant negative correlation between the structural variation and optimization of vowel systems (Spearman’s *rho* = -0.3474, *p*-value = 1.7419×10^-16^). This suggests that a vowel system with low variation tends to rearrange its structure to reach optimal; however, when a vowel system almost reaches an optimal one, its optimization will decrease since there is not much space to adapt the structure. This dynamic process echoes the self-adaptation of vowel systems.

### Inter- and intra-regional differences of the structural variation and optimization of vowel systems

For inter-regional comparison, we performed a one-way analysis of variance (ANOVA) on the structural variation and optimization of vowel systems across eight geographic regions (Fig. 1a-b). The analysis reported significant inter-regional differences in the structural variation and optimization of vowel systems (as for variation: *p*-value < 1.2876× 10^-7^; as for optimization: *p*-value < 5.7794× 10^-14^). Among the median values of structural variation, East Asia had the maximum median value and Middle East had the minimum value (see Fig. 1c). Among the median values of structural optimization, Africa had the minimum value and North-Central America had the maximum value (see Fig. 1d). In addition, Fig 1e shows that the median values of structural variation were negatively correlated with the median values of structural optimization across all eight geographic regions (Spearman’s *rho* = -0.8095, *p*-value = 0.0218). These findings indicate significant inter-regional differences of vocalic structures in the world.

**Figure. 1.**
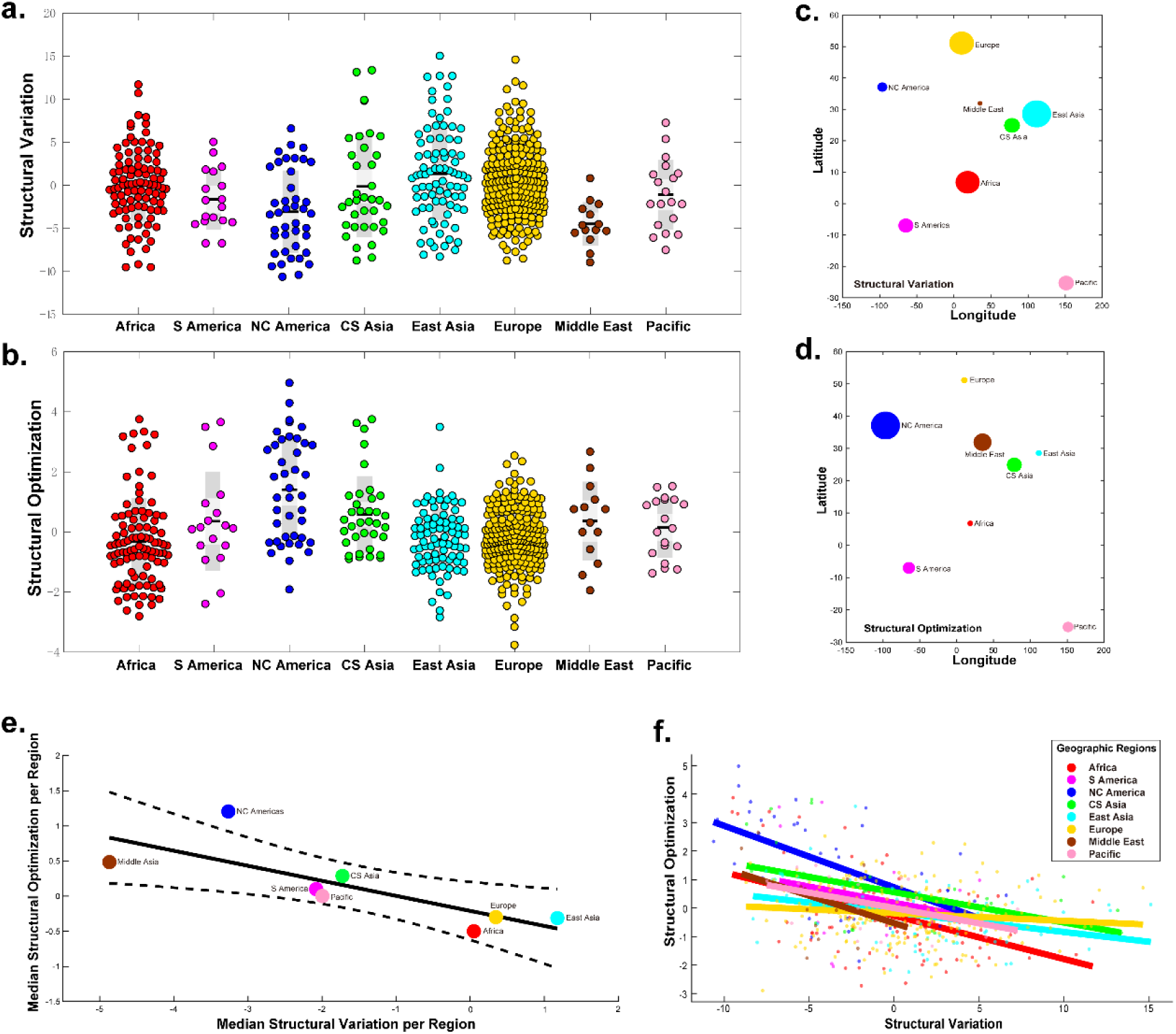
Inter- and intra-regional differences for structural properties of vowel systems. The structural variation (a) and optimization (b) in different geographic regions are shown in the Univar scatter plots. The median structural variation (c) and optimization (d) are displayed in each geographic region. Circle size corresponds to values of these two structural properties. The relation of these two indices in the eight geographic regions is examined by a linear regression model (e). The dotted lines represent 95% Confidence Intervals. The linear fits (colored lines) in (f) reveal a consistent negative relationship between structural variation and optimization across all geographic regions. Points are color coded according to different regions. NC America: North-Central America; S America: South America; CS Asia: Central-South Asia.

Intra-regional correlations between the structural variation and optimization of vowel systems in all eight regions were negative (Table 1). However, correlations in South America, Middle East and Pacific were not significant, probably due to the small sample sizes. Nonetheless, these findings suggest that the structural variation and optimization of vowel systems are consistently negatively correlated in different geographical regions; in other words, such tendencies could be universal in the world.

**Table 1.**
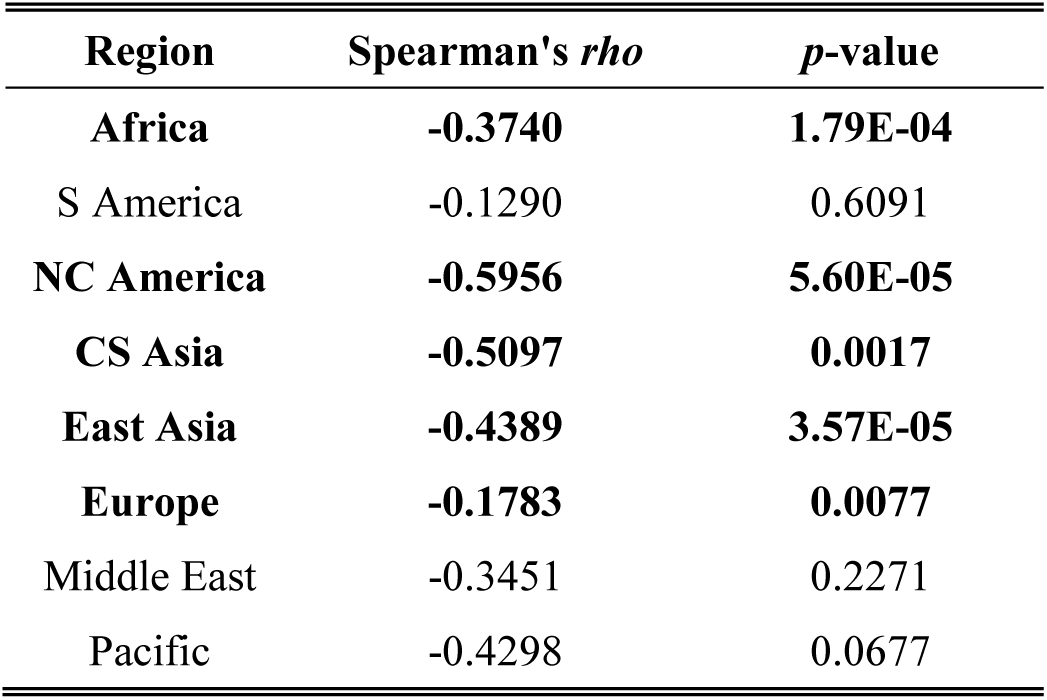
The results of Spearman’s correlation between structural variation and optimization values of language samples in each geographic region. Significant correlations are marked in bold.

| **Region** | **Spearman's $\rho$** | **$p$-value** |
| --- | --- | --- |
| **Africa** | **-0.3740** | **1.79E-04** |
| S America | -0.1290 | 0.6091 |
| **NC America** | **-0.5956** | **5.60E-05** |
| **CS Asia** | **-0.5097** | **0.0017** |
| **East Asia** | **-0.4389** | **3.57E-05** |
| **Europe** | **-0.1783** | **0.0077** |
| Middle East | -0.3451 | 0.2271 |
| Pacific | -0.4298 | 0.0677 |

### Coevolution between the structural variation and optimization of vowel systems

The above synchronic analyses reveal a universal negative correlation between the structural variation and optimization of vowel systems. To further address how these two properties of vowel systems evolved in history, we applied phylogenetic comparative methods on a family tree of Indo-European (IE) languages from (35). This was a rooted tree established based on the 200 Swadesh lexical cognates. We focused on the 33 contemporary IE languages appearing in both datasets (see Fig. 2). The phylogenetic comparative analysis revealed decisively a strong negatively correlated evolution between the structural variation and optimization of vowel systems (see Table 2). This is in line with our conclusion in the synchronic comparisons: The evolution of vowel systems is a dynamic process of self-adaptive rearrangement of their structure and optimization capacities, and such process is reasonably universal across languages.

**Figure. 2.**
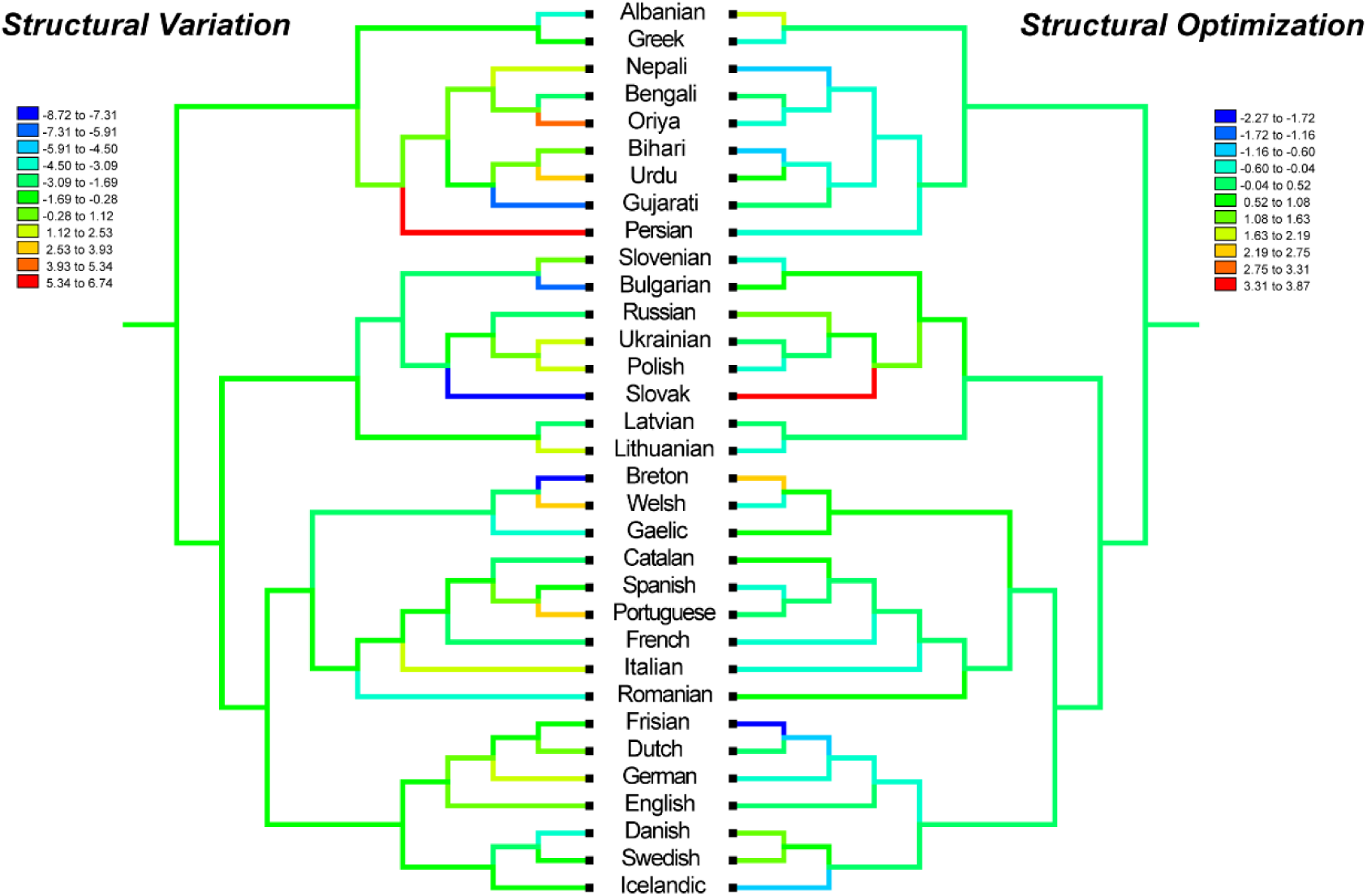
The mirror phylogenetic trees of 33 contemporary Indo-European languages with structural variation and optimization values. The basic tree model is drawn from (35). The ancestral states of each observed indices are reconstructed based on Parsimony method to trace the character history, implemented in Mesquite. Branch colors represent estimated ancestral values of two structural properties.

**Table 2.** . Model fit statistics and fixed-effects estimates for predicting the three non-linguistic outcomes (latitude, longitude, and log-transformed speaker population size) (*N* = 532). Shades in grey highlight the absolute *t*-statistic for fixed-effects greater than 2.

| Outcome | Model fit statistics |  |  | Predictor | Fixed Effects |  |  |  | Tag |
| --- | --- | --- | --- | --- | --- | --- | --- | --- | --- |
|  | AIC | BIC | Log-Likelihood |  | Coefficient | SE | t-statistic | p-value |  |
| Latitude | 4866.7 | 4888.1 | -2428.4 | Intercept | 33.557 | 3.4061 | 9.8519 | 3.8602E-21 |  |
|  |  |  |  | Structural Variation | 0.099862 | 0.22857 | 0.43689 | 0.66237 |  |
|  | 4866.8 | 4888.1 | -2428.4 | Intercept | 33.534 | 3.3828 | 9.9129 | 2.3078E-21 |  |
|  |  |  |  | Structural Optimization | 0.2921 | 0.796 | 0.36696 | 0.71379 |  |
|  | 4863.2 | 4884.5 | -2426.6 | Intercept | 26.887 | 4.8284 | 5.5685 | 4.09E-08 |  |
|  |  |  |  | Vowels | 0.93917 | 0.48506 | 1.9362 | 0.053373 |  |
| Longitude | 5899.4 | 5920.8 | -2944.7 | Intercept | 28.047 | 3.212 | 8.732 | 3.2748E-17 |  |
|  |  |  |  | Structural Variation | 2.0172 | 0.57351 | 3.5172 | 0.0004735 | *** |
|  | 5889.4 | 5910.7 | -2939.7 | Intercept | 28.424 | 3.8448 | 7.3929 | 5.6393E-13 |  |
|  |  |  |  | Structural Optimization | -10.039 | 2.0671 | -4.8565 | 1.5754E-06 | *** |
|  | 5900.9 | 5922.3 | -2945.4 | Intercept | 0.038983 | 8.9099 | 0.004375 | 0.99651 |  |
|  |  |  |  | Vowels | 4.066 | 1.2576 | 3.2332 | 0.0013005 | ** |
| Log10(Speaker population size) | 2003.7 | 2025.1 | -996.84 | Intercept | 6.3401 | 0.27949 | 22.685 | 4.015E-80 |  |
|  |  |  |  | Structural Variation | 0.03166 | 0.015588 | 2.0311 | 0.042744 | * |
|  | 2003.1 | 2024.5 | -996.54 | Intercept | 6.3162 | 0.29439 | 21.455 | 5.6976E-74 |  |
|  |  |  |  | Structural Optimization | -0.11649 | 0.053894 | -2.1614 | 0.031108 | * |
|  | 2005.5 | 2026.9 | -997.74 | Intercept | 5.9664 | 0.36825 | 16.202 | 2.982E-48 |  |
|  |  |  |  | Vowels | 0.049687 | 0.033063 | 1.5028 | 0.13349 |  |

Note: \*: $p$ -value < 0.05; \*\*: $p$ -value < 0.01; \*\*\*: $p$ -value < 0.001.

We also examined the evolutionary mode and tempo of the two structural properties. Estimates of Pagel’s *λ* showed that both variation and optimization exhibited language-independent evolutions with *λ* equivalent to 0 (see Table 3); in other words, each vowel system had its unique evolutionary process. Estimates of the *κ* parameter indicated that the structural variation primarily followed a gradual evolution with increased rates of evolution in shorter branches, but stasis in longer ones (0 < *κ* < 1, see Table 3). By contrast, the evolution of the structural optimization had a punctuated mode, indicating that the optimization of a vowel system could change in burst mode, followed by a long time stasis. Estimates of the *δ* parameter (significantly larger than 1) revealed that the evolution of vowel systems was a process of languages-specific adaptation.

**Table 3.** . Likelihoods for dependent and independent models of correlated evolution between structural variation and optimization.

|  | **Log Likelihood** | **STD** | **SE** | **Correlation** | **STD** | **Log<sub>10</sub> Bayes factor** |
| --- | --- | --- | --- | --- | --- | --- |
| Dependent model | -148.00 | ±1.07 | ±0.03 | -0.54 | ±0.02 | 11.11 |
| Independent model | -153.56 | ±1.02 | ±0.03 | 0.00 | - |  |

Note: The log<sub>10</sub> Bayes factor (Log<sub>10</sub> BF) indicates the relative support for the dependent model over the independent model. Values < 2 suggest weak support, > 2 positive support, 5-10 strong support, and > 10 decisive support for the dependent over the independent model. Log likelihood for each model is a marginal.

### Correlations between the structural properties of vowel systems and the non-linguistic factors

Regarding the controversial relationship between non-linguistic factors (e.g., speaker population size, geographic coordinates) and vowel inventory, we established linear mixed-effects models with random intercept to examine the correlations between different structural properties of vowel systems and the two non-linguistic factors (see Materials and Methods).

Statistical results showed that neither structural variation nor optimization were correlated with the latitudinal distribution of all language samples, but both were significantly correlated with the speaker population sizes and the longitudinal distribution of languages (see Table 4). The coefficient of the structural variation for predicting population size was positive, but that for predicting population size was negative. These indicate that a language with a larger speaker population size tends to self-adaptively rearrange its vowel system to reach an optimal structure; when the structure of a language is closer to its optimal state, its optimization capacity may diminish. Similarly, a language with a larger longitude value tends to have a higher structural variation and a weaker optimization capacity. These results reveal the significant roles of speaker population size and longitudinal location in shaping the vocalic structures of languages. In addition, the vowel inventory size was significantly correlated with the longitude of language, but not latitude nor speaker population size. This implies a serial founder effect of vowel systems in the world (36, 37).

**Table 4.** Log_10_ BF tests for the observed versus expected values of phylogenetic scaling parameters for different models of trait evolution of vowel systems. The selected models are shown in bold.

| Trait | Observed value | Log likelihood | Log10 BF |
| --- | --- | --- | --- |
| **Structural Variation** |  |  |  |
| Lambda $\lambda$ | | | |
| $\lambda$ estimated | 0.1544 | -88.7514 | |
| $\lambda$ forced = 1 | | -96.2761 | 15.0494 |
| **$\lambda$ forced = 0** |  | **-88.5911** | **-0.3206** |
| Kappa $\kappa$ | | | |
| **$\kappa$ estimated** | **0.6572** | **-95.3131** |  |
| $\kappa$ forced = 1 | | -96.4627 | 2.2992 |
| $\kappa$ forced = 0 | | -96.4041 | 2.1820 |
| Delta $\delta$ | | | |
| **$\delta$ estimated** | **2.9802** | **-94.3859** |  |
| $\delta$ forced = 1 | | -97.6510 | 6.5302 |
| **Structural Optimization** |  |  |  |
| Lambda $\lambda$ | | | |
| $\lambda$ estimated | 0.2075 | -50.9231 | |
| $\lambda$ forced = 1 | | -56.9023 | 11.9584 |
| **$\lambda$ forced = 0** |  | **-48.9387** | **-3.9688** |
| Kappa $\kappa$ | | | |
| $\kappa$ estimated | 0.2319 | -55.8840 | |
| $\kappa$ forced = 1 | | -58.7054 | 5.6428 |
| **$\kappa$ forced = 0** |  | **-55.5030** | **-0.7620** |
| Delta $\delta$ | | | |
| **$\delta$ estimated** | **2.5358** | **-53.2956** |  |
| $\delta$ forced = 1 | | -57.2178 | 7.8444 |

Note: The absolute log<sub>10</sub> BF > 2 represents the significance, and the negative value supports the latter model.
Log Likelihood is actually Log Marginal Likelihood.
The observed parameters ( $\lambda$ , $\kappa$ , $\delta$ ) were contrasted with values expected under the null hypothesis (value = 0 and 1). When the observed models show no significant difference from expected models, the latter was selected.

## Discussion

To capture various optimization capacities of the whole vowel system, this paper extended the concept of optimization from perceptual domain into articulatory reality. We obtained the composite parameters of structural variation and optimization of vowel systems from dispersion, focalization, articulatory space, and their corresponding optimization capacities. Synchronic comparison of these two properties revealed a universal negative correlation between them in world languages. This tendency was also ascertained as a correlated evolution in the diachronic analysis based on 33 contemporary Indo-European languages, though the evolution of structural variation appears to be gradual and that of structural optimization appears to be punctuated. Among non-linguistic factors, our analyses confirmed that the structural differences of vowel systems were correlated with the longitudes and speaker population sizes of languages.

Evolution of human language is a process of gradual optimization in both articulatory and auditory domains. The universal negative correlation between the structural variation and optimization of vowel systems accounts for this process: The more variations a vowel system has, the low optimization it exhibits; in other words, the closer a vowel system is to its own optimal structure, the fewer space it has for improvement. Rather than saying that all vowel systems reach the same optimal system, we stress that each system is moving toward its own optimal state, respectively. The variation-optimization correlation could be due to articulatory limitations, such as the opening and closing ranges of jaw or mouth, or tongue activity space. The negativity of the correlation was observed both inter- and intra-regionally, though not all intra-regional linear models showed significance. This could result from specific cultural and demographic activities in some local regions.

Relationship between phonological inventory and speaker population size of languages is under a hot debate. Our study showed that structural changes of vowel systems were significantly correlated with speaker population sizes. With growth in population size, speech systems start to show more individual variations, which could cause more ambiguities and reduced communicative effectiveness. This obviously conflicts with the primary functional goal of language. As a consequence, language has to adapt its own system to fulfil the communicative goal. Taking vowel systems as example, expanding pairwise distances between vowels can obtain a wider range of vowel variations in the perceptual domain, and speaking in exaggerated pronunciation manners can avoid confusion in speech production (15). These explain why speaker population size, rather than vowel inventory size, is correlated with structural properties of vowel systems. Such self-rearrangement is not only universal in speech, but also evident in other language systems, such as the dependency length minimization in syntax (1, 2) and the simplification in morphology (38, 39).

There lacks explicit evidence that speaker population size could explain a significant proportion of variation in vowel inventory size (29, 30). Our study reported no global correlation between them, in line with the conclusions of (28). However, we could not totally exclude the possibilities of local correlation between the two in some regions, especially multilingual regions (e.g., colonies). The drastic social structures in these regions are commonly accompanied with deep language contact or admixture. A typical process as such is language transfer, and many phonological changes could occur along with this process, such as under-differentiation (merge of two sounds that are distinct in the second language but not in the first), over-differentiation (imposition of phonemic distinctions from the first language on allophones of a single phoneme in the second), or out-right substitution (using a phoneme from the first language for a similar but distinct phoneme in the second) (40, 41). Vowel inventory size is directly increased or decreased in the first two cases, but remain unchanged in the third case. All these could eventually result in a weak correlation between population size and vowel inventory size. Note that this is just one possibility out of many.

Our analysis identified significant correlations between structural differences of vowel systems and longitude coordinates of languages. This implies a serial founder effect similar to Atkinson’s conclusion (23), but we do not intend to support the hypothesis of serial founder effect in language evolution, because compared to genes, sounds are more prone to borrow during contact and environmental factors also play a role in linguistic structure during language evolution (32, 42).

Synchronic linguistic typology is a reflex of diachronic change (7). Traditional typological investigations are based primarily on correlation analysis, which is insufficient to reconstruct the evolutionary histories of languages (43). We applied phylogenetic analysis to quantify the process and mode of the evolution of linguistic features, which are critical to understand many questions in historical linguistics (44). Our phylogenetic analyses based on the Indo-European languages provided strong evidence of language-specific adaptations in the evolution of vowel systems (Fig. 2): The structural variation of vowel systems proceeds in a gradual mode, whereas the optimization proceeds in a punctuated mode.

The gradual mode indicates that languages accumulate over time inconspicuous changes in typological structures of vowel systems, such as numerous acoustic variations of individual vowels in a speaker population (45, 46). Vowel systems tend to evolve so as to achieve efficient speech understanding and high speech intelligibility under a variety of conditions and disturbances (47). As an analogy to biological evolution, this continuous fine-tuning process could be induced by social selection. In contrast, the punctuated mode of the optimization of vowel systems implies that languages rearrange their own vowel systems rather rapidly, and maintain the new structures for a relatively long time. There are two mechanisms that could drive for such mode of evolution. The first one is the founder effects, which could explain the structural rearrangements occurring during language origin and divergence. The second one is language contact or admixture induced by demographic activities, which could invoke structural changes as one language enters a new linguistic environment. Such changes usually manifest during second language acquisition in local population or immigrants. Such punctuated mode was identified not only in phonology, but also in other language systems such as lexicon (48). The phylogenetic analysis of the structural variation and optimization of vowel systems provides explicit evidence that languages slowly accumulate structural changes until a threshold is reached, and then the whole system is rapidly restructured (49). The inconsistent modes of the two properties of vowel systems reflect distinct yet correlated evolutionary processes, which ask for further investigations based on empirical data of other language families.

## Materials and Methods

### Formant data processing based on Becker’s vowel inventory corpus

This paper used formant frequencies of vowels in different languages for analysis. The data were all taken from Becker-Kristal’s vowel inventory corpus (50). The corpus contains in total 555 samples of 357 languages and/or dialects. The formant frequencies of the vowels in the samples were extracted from recorded raw data by five elicitation methods: stained isolated vowels, vowels embedded in isolated word lists, vowels embedded in words placed in carrier sentences, vowels embedded in words placed in meaningful sentences, and vowels pronounced as parts of speech flow such as conversations. The vowel inventories of the languages were classified into six quantity-contrastive constituents: combined, mixed, long, short, uniform and unknown. After filtering out samples having the incomplete first two formant frequencies of vowels, we obtained 532 samples covering 357 languages and/or dialects for the proposed analyses. All these language samples have valid geographic coordinates (latitudes and longitudes) and speaker population sizes that match Ruhlen’s database (30), the PHOIBLE database(8), and the AUtotYP database (51). The geographical locations of these samples cover in total eight world regions: Africa (Sample size N = 97), Europe (N = 223), Middle East (N=14), Central South Asia (CS Asia; N = 36), East Asia (N = 84), Pacific (N = 19), South America (S America; N = 18) and North Central America (NC America; N = 41) (see Table S1).

### Psycho-acoustic conversion of formant data

We transferred the formant frequencies of vowels into a psychoacoustic scale. This conversion is necessary because the resolution of the human auditory system is determined by the critical band analysis following a non-linear Bark scale, such that high frequency sounds appear closer together than low frequency sounds (52). In practice, the use of the Bark scale stretches the vowel space where human ears are most sensitive, and contracts the space where frequency differences are difficult to perceive by human ears (53). For example, human ears have a high discrimination sensitivity in the low frequency domain (e.g., from 500 Hz to 1000 Hz), but a weak sensitivity in the high frequency domain (e.g., from 4500 Hz to 5000 Hz). The auditory Bark scale (53) that we used is defined as in equation (1):

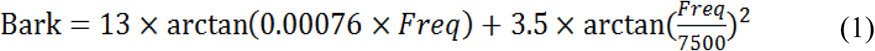

### Dispersion estimates

Dispersion is one systematic properties of vowel system. Dispersion (DE) accumulates the reciprocal of total perceptual distances between each pair of vowels in a vowel system. It can be used to predict the most frequent vowel system(s) among languages and identify the most optimal one(s) in terms of clarity. A large DE of a vowel system indicates that the vowels therein are crowded in the vowel space (54). In this paper, we modified the original equation for DE estimates proposed in (10). Given a system of *n* vowels, DE was measured as in equation (2):
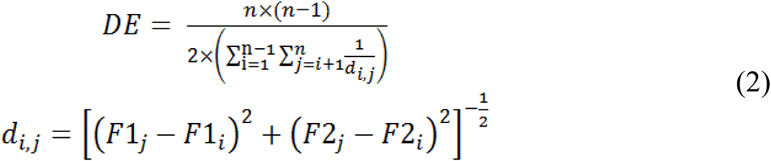

where F1*_i_*, F2*_i_*, F1*_j_*, F2*_j_* are the Bark values of the first and second formants of vowels *i* and *j*. Simply put, DE is the harmonic average of the inverse-squared Euclidean distances between each possible pair of vowels in a vowel system. If vowels are very close to each other in a system, DE of this system is small.

### Focalization estimates

Focalization (FE) is another systematic properties of vowel system (11). Focalization measures the relative perceptual distance between adjacent formants of vowels, which assigns a focal quality to each vowel. They determine whether or not the vowel configuration produced by articulatory manoeuvres is stable (54). Vowels with larger FE can be perceived more explicitly. FE was measured as in equation (3):
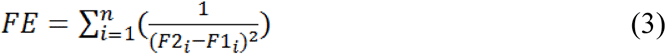

FE of each vowel is the sum of the inverse-squares of the adjacent (F1 and F2) formant difference of the vowel in the system. The most focal vowels in our dataset are /u/ and /ɯ/ because their first two formants are much closer than those of the others.

### Articulatory space estimates

Articulatory space (AS) for a vowel system is an alternative systematic property. A traditional method of estimating articulatory space is based on the Euclidean space formed by corner vowels. This method is not sufficiently sensitive to production deficits. In this paper, we adopted the Convex-Hull approach (55) in computational geometry to calculate the articulatory space areas of the vowel systems in different languages. Mathematically speaking, a convex hull of a set of X points in a Euclidean space is the smallest convex set that contains X (56). Convex-hull estimation depends on the relative positions of the vowels in a vowel space, which is more efficient and accurate for estimating the articulatory space area than the traditional method. We used the function *convhull* in Matlab R2015b to calculate the convex hull of each vowel system.

### Assessing optimization of vowel systems using Monte Carlo Sampling

In order to compare the real vowel system with the random ones, we conducted an analysis of optimality using Monte Carlo Sampling, a method similar to the one proposed in (34). It is common that no formant pairs have their centres separated by less than 1 Bark (57). Thus, we added a basic perceptual constraint in the sampling approach that the adjacency of two formants was beyond 1 bark. The constraints integrated into the basic sampling process as follows:

1. For each language sample, obtain the largest and smallest values of the first and second formants (F1_max_, F1_min_, F2_max_, F2_min_) of the vowels therein. These values define the boundaries of the vowel space in the sample.
2. Generate a single random vowel by selecting its F1 uniformly within the range [F1_min_, F1_max_] and its F2 uniformly within the range [max(F1, F2_min_), F2_max_], to ensure a smaller F1 than F2. If the difference between F1 and F2 is smaller than 1 Bark scale, reselect these values.
3. Repeat Step 2 to generate the same number of random vowels as that of the natural vowels in the language sample.
4. Repeat Steps 2 and 3 to create 10,000 sets of random vowel systems.
5. Calculate the standard scores (*z*-scores) based on dispersion (DE), focalization (FE) and articulatory space (AS) between the real vowel system and the 10,000 random sets, as in equation (4):
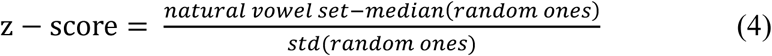

The *z*-scores for dispersion, focalization and articulatory space indicate the capacities of systematic optimization at a global level. An absolute *z*-score greater than 0.0 indicates that the vowels in a language sample are further apart from one another than expected by chance. We calculated three optimization values (*z*-scores) for dispersion, focalization, and articulatory space of each vowel system.

### Structural variation and optimization of vowel systems

We applied Principal Component Analysis (PCA) on the three systematic properties (dispersion, focalization and articulatory space) to obtain the structural variation of vowel systems. Similarly, the structural optimization of vowel systems was obtained via applying PCA on the three corresponding optimization values. This approach was different from (11), in which they used a simple the weighted summation function. PCA was implemented in Matlab 2015b.

### Linear mixed-effect regression analyses

We used linear mixed-effects regression models to examine correlations between structural properties of vowel systems and non-linguistic factors. Mixed-effects regression models allow for simultaneous consideration of multiple covariates, and keep the between-participants and between-items variance under statistical control (58, 59). In each model, the vowel size, structural variation and optimization were treated as independent variables (X), or fixed effects. Each model also included two intercepts respectively for the elicitation methods in recording and the quantity-contrastive constituents of vowel inventories, which were treated as group-level random effects that could affect the intercepts of regressions. Although maximal random effect structures including random slopes are theoretically desirable (60), such complicated models were not pursued here in consideration of practical constraints on model convergence.

We fit three models targeting respectively three dependent variables (Y): latitude, longitude, and log-transformed speaker population size (for multiplicative scale). The linear mixed-effects models have the format as in equation (5):
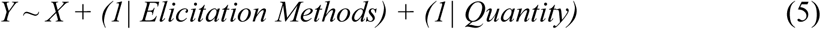

### Modelling trait evolution in vowel systems

We used a random-walk Markov chain Monte Carlo (MCMC) procedure in BayesTraits-Continuous to examine separate evolution of individual continuous trait and correlated evolution of two traits over phylogeny. The basic phylogenetic tree used here was a published Bayesian cognate-based linguistic tree of 33 contemporary Indo-European (IE) language (35). This was a dated and rooted tree. In order to obtain insights on the evolutionary history of the vowel systems of the IE languages, we reconstructed the ancestral structural properties and traced the evolutionary histories of them on the IE language tree. The reconstruction was on the basis of parsimony method implemented in MESQUITE 3.3.1 (61). Here, the continuous traits were structure variation and optimization of vowel systems. We used 100 stepping stones and 1000 iterations per stone to estimate the marginal likelihood. Total iteration was set as 1,010,000 and the first 10,000 was Burn-in.

Pagel’s lambda (*λ*), kappa (*κ*) and delta (*δ*) were estimated based on the phylogeny of the IE languages to determine respectively the phylogenetic associations, mode and tempo of trait evolution. These three parameters were all calculated by BayesTraits V3 program (62).

The parameter *λ* assesses the phylogenetic contribution to a given trait. *λ* = 0 indicates that the trait evolution has proceeded independently of phylogeny, whereas *λ* = 1 suggests that the trait evolves as expected given the tree topology and the random walk model (i.e., Brownian motion). Intermediate values of 0 < *λ* < 1 indicate different degrees of the phylogenetic signal.

The branch-length scaling *κ* is used to test for a punctuated versus a gradual mode of trait evolution in phylogeny. *κ* < 1 indicates proportionally more evolution in shorter branches, whereas *κ* > 1 suggests proportionally more evolution in longer branches. The latter one is consistent with a gradual mode of evolution. The extreme of *κ* = 0 suggests a punctuated mode of evolution.

The path scaling parameter *δ* detects whether the rate of trait evolution has accelerated or slowed over time in a given phylogenetic tree. It can find evidence for: adaptive radiations (*δ* < 1), temporally early trait evolution or early burst; gradual evolution (*δ* = 1) with a constant rate; or specific adaption (*δ* > 1), temporally later trait evolution with acceleration over time.

We calculated the log-transformed (base 10) Bayes factor to compare model fits. This factor can illustrate the weight of evidence in support of one model over another. A value between 0 and 2 indicates weak evidence, and over 2 positive evidence. A value between 5 and 10 indicates strong evidence, and over 10 very strong evidence.

